# Identifying likely transmission pairs with pathogen sequence data using Kolmogorov Forward Equations; an application to *M.bovis* in cattle and badgers

**DOI:** 10.1101/2020.06.11.146894

**Authors:** Gianluigi Rossi, Joseph Crispell, Daniel Balaz, Samantha J. Lycett, Richard J. Delahay, Rowland R. Kao

## Abstract

Established methods for whole-genome-sequencing (WGS) technology allow for the detection of single-nucleotide polymorphisms (SNPs) in the pathogen genomes sourced from host samples. The information obtained can be used to track the pathogen’s evolution in time and potentially identify ‘who-infected-whom’ with unprecedented accuracy. Successful methods include ‘phylodynamic approaches’ that integrate evolutionary and epidemiological data. However, they are typically computationally intensive, require extensive data, and are best applied when there is a strong molecular clock signal and substantial pathogen diversity.

To determine how much transmission information can be inferred when pathogen genetic diversity is low and metadata limited, we propose an analytical approach that combines pathogen WGS data and sampling times from infected hosts. It accounts for ‘between-scale’ processes, in particular within-host pathogen evolution and between-host transmission. We applied this to a well-characterised population with an endemic *Mycobacterium bovis* (the causative agent of bovine/zoonotic tuberculosis, bTB) infection.

Our results show that, even with such limited data and low diversity, the computation of the transmission probability between host pairs can help discriminate between likely and unlikely infection pathways and therefore help to identify potential transmission networks, but can be sensitive to assumptions about within-host evolution.

## 1. Introduction

In recent years, network models have been increasingly used to represent the complex set of interactions (i.e. contacts) that can lead to pathogen transmission in humans ^1,2^, wildlife ^3^, and livestock ^4,5^. In a network paradigm, individuals or groups of hosts (i.e. farms, social groups, or sub-populations) are represented as *nodes* (in graph theory, *vertices*), and the potential infectious contacts between them as *links (edges)*.

An important distinction exists between the *contact network* and the *transmission network*: while the former includes all potential transmission contacts, the latter is a subset of the former describing pathogen transmission patterns ^5,6^. Identifying the transmission network, even when the contact network is well described, can be a challenging filtering process informed by multiple factors. Techniques are still needed to disentangle these factors, using the different sources of information (evolutionary, immunological, and epidemiological) available to infer likely transmission pathways. Most importantly, we wish to know “how likely is it that individual A infected individual B?”, or “how likely is it that a third unsampled individual was involved in the transmission chain between individuals A and B?”, the key questions in *forensic* or *‘precision’ epidemiology*. The answers to these questions, and transmission pathway reconstruction, are important for gathering information about outbreaks, to shed light on transmission dynamics, and to help infer epidemiological parameters.

Whole genome sequencing (WGS) can be used to detect polymorphisms in a genome with high resolution, and therefore discriminate between closely related strains. Polymorphisms are caused by errors that occur during pathogen replication within the host. Generally these single nucleotide polymorphisms (SNPs) are considered to be neutral in bacterial species within the timescale of disease outbreaks ^7^. In the absence of horizontal genetic transfer, tracking these SNPs would be expected to follow the pattern of transmission. In combination with an increasing ability to extract genetic material (either directly from clinical samples or from cultured isolates) and with rapid and minimal processing, large-scale characterization of populations of pathogen genomes is now possible ^8–10^. These advances have proven to be transformative for forensic epidemiology, especially when populations can be sampled densely.

The observed genome diversity in a population of pathogens is the result of processes happening at two different scales. The two most important of these are the evolution of the pathogen’s genome within the host and its transmission to another host ^11^. Both processes are subject to population bottlenecks that could limit strain circulation both within and between hosts ^12^. At larger scales, pathogen genotype patterns may be influenced by the contact network; i.e. host social organisation and movement behaviour will determine contact rates within and between host populations. Contact patterns are especially important where disease prevalence and cross-immunity between strains are both high^13^, as this leads to substantial transmission-blocking due to prior infection (and therefore alteration of the effective transmission network by the history of the pathogen itself). Given the availability of genomic data, the most straightforward approach to describe the relationship between hosts would be based on the genetic clustering of the sampled pathogen strains. However, this would not provide any information about the direction of transmission ^11^. Direction can be estimated by coalescent-based phylodynamic models that infer disease dynamics from an observed genealogy but with only indirect reference to the underlying host demographics ^14^. Phylodynamic approaches have been extended further to include different stages of infection and structured host populations ^15^, and later an underlying contact network ^16^. In the latter case a pairwise coalescent model was embedded within an individual-based stochastic simulation model; this analysis showed that the host contact network can interact with the timing of coalescent events during an epidemic.

Other approaches have considered likelihood-based frameworks, where the likelihood function defines the probability of a transmission tree given temporal and genetic information ^17,18^. These likelihood-based functions can then be used to estimate the probability of a transmission tree and the parameters of the function itself. In addition, the likelihood framework can account for missing data. This framework approach has been extended further ^19^, to show the influence of pathogen within-host dynamics on the relationship between transmission and the phylogenetic tree. A framework based on Bayesian inference embedding an underlying mathematical model developed by Morelli et al.^20^ considered the geographical distance in the transmission probability definition. Another Bayesian inference method ^21^ was used to infer transmission pathways considering the evolutionary and epidemiological processes simultaneously but did not consider the within-host dynamics of the pathogen, assuming instead that a single dominant strain propagates within and between clusters of hosts. Similarly, a structured-coalescent evolutionary model (SCOTTI) ^22^ took a Bayesian approach to reconstruct the transmission events within outbreaks. The SCOTTI framework represents the transmission process as migration events between *populations of pathogens* (i.e. hosts). Li et al. ^23^ used a similar method based on particle Markov Chain Monte Carlo (MCMC) to estimate the transmission heterogeneity (i.e. offspring distribution) from incidence time series and pathogen phylogeny. Maximum parsimony algorithms have also been used to estimate transmission events, by minimising the number of infections consistent with the identified ancestral states in the tree. Romero-Severson et al.^24^ used this criterion with a coalescent HIV model to evaluate the transmission histories of two hosts; they showed that the direction of transmission and the presence of an unsampled intermediary or a common source could be included or excluded depending on the relationship between the two strains and the number of lineages transmitted. The parsimony criteria was also used by Wymant et al. ^11^ to develop Phyloscanner, a software tool that can determine transmission pathways from multiple genotypes per infected host. These approaches have produced extremely powerful tools to help disentangle the relationship between pathogen evolution and the infection processes^25^. Many of these have been successfully applied to rapidly evolving RNA viruses such as HIV, Ebola, influenza or foot-and-mouth disease. On the other hand however, Campbell et al. ^26^ showed that sequence data for pathogens with lower *transmission divergence* (defined as the number of mutations separating whole genome sequences sampled from transmission host pairs) provide little information about individual transmission events on their own. Hence, estimating transmission direction is particularly difficult for pathogens such as *Mycobacterium bovis* (a clonal pathogen that is a causative agent of bovine/zoonotic/animal tuberculosis or bTB). *M. bovis* is characterized by an extremely slow and highly variable substitution rate, generating low and uncertain levels of genetic diversity, especially when considering small clusters of closely related infections ^27–29^. At this scale *M. bovis* is expected to experience no horizontal genetic transfer, and therefore there is a close correspondence between the phylogenetic tree and the transmission network; nevertheless identifying transmission chains is particularly challenging. A further consideration is that the majority of methods require dense and/or extensive metadata to infer transmission patterns, which are often not available. This is often true for pathogens where wildlife are involved, as data on wildlife populations can be both sparse and imprecise.

Here, we investigated the discriminatory power of sequence data to identify transmission pathways where SNPs are rare, genetic diversity per transmission generation is low and highly variable, available sequences may be few (here, considering pairs or triplets of sequences), and metadata is limited to the recorded sampling times, as is often the case when data are convenience sampled in livestock and wildlife. To maximise the information from such data, we adopt a probabilistic approach to capture both within-host pathogen evolution (i.e. new SNP substitutions) and between-host transmission using the Kolmogorov Forward Equations (KFEs). Instead of recording the number (or fraction) of individuals in a given infection state (e.g. number of Susceptible, Infectious and Recovered in the SIR model case), the KFEs describe the probability of the system having a given state, with an exact number of individuals in each infected state ^30–32^. We proceeded in a pairwise fashion, thus the system status was defined by the combination of infection states of two hosts and the SNPs differences in the pathogen strains they are infected with. We tested this method on a simulated transmission tree and on a bTB infected population of badger and cattle in Woodchester Park (England). We used this method to test contrasting model assumptions, as well as to assess the likely epidemiological importance of different contact mechanisms.

## 2. Results

### 2.1 Woodchester Park cattle and badger population

In the present analysis we used the Kolmogorov Forward Equations to calculate the direct transmission probability amongst badgers, amongst cattle, and between the two species. We follow the approach taken by Sharkey ^31^, who used the KFEs to describe infection dynamics at the individual and pairwise level, but here adding in states to describe the evolution of the pathogen. We applied the pairwise KFE model to data from an endemic *M. bovis* population circulating in cattle (*Bos taurus*) and European badgers (*Meles meles*) in Woodchester Park, Gloucestershire (UK). Since 1977, the population of badgers residing in the Woodchester Park study area has been the subject of an ongoing capture-mark-recapture project ^33^. Following an earlier phylogenetic analysis by Crispell et al.^34^, the infected population was divided in five clades according to their genetic distance. These clades (subclades in the case of clade 4) were used as proxies for network clusters, as we considered clade members to be potentially in contact with one another. The transmission probability varied substantially between pairs (median[95^th^ quantile] l 0.26 × 10^−4^ [0 – 0.03]) and was negatively correlated with SNP distance (Fig 1). Although all five clades included strains that have been considered to be closely related (< 20 SNPs distance, ^29^), even a small SNP difference (up to five) has been shown to be useful in discriminating differences in the likelihood of a transmission event between two sampled individuals.

**Fig 1.**
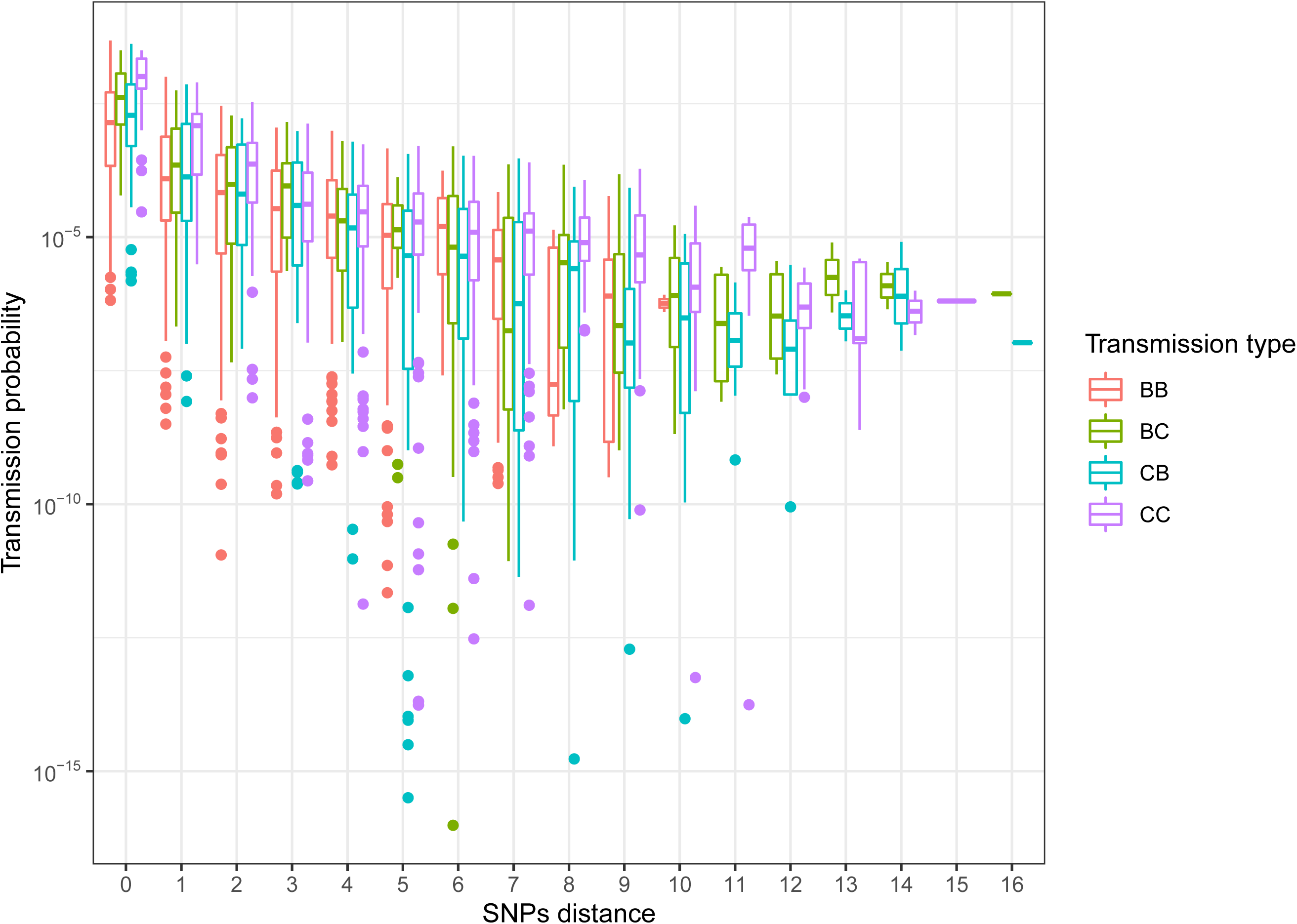
Pairwise transmission probability. The pairwise transmission probability (y-axes) for each host pair vs. the SNP distance (x-axis).

While data on geographic distances between individuals and group (social groups for badgers, herds for cattle) affiliations were available, we deliberately excluded them from the analysis to determine if the effect of distance could be partially recovered by genetic and sample time data alone. As shown in Figure 2, the proposed method captured differences in the likelihood of transmission between pairs of hosts regardless of whether they belonged to the same social group (i.e. farm or sett, panel A) and species (panel B), although we observed a substantial overlap in the distributions. On the other hand, at the spatial scale considered here (≤ 10km) the distance in space did not seem to affect the transmission probability (panel D).

**Fig 2.**
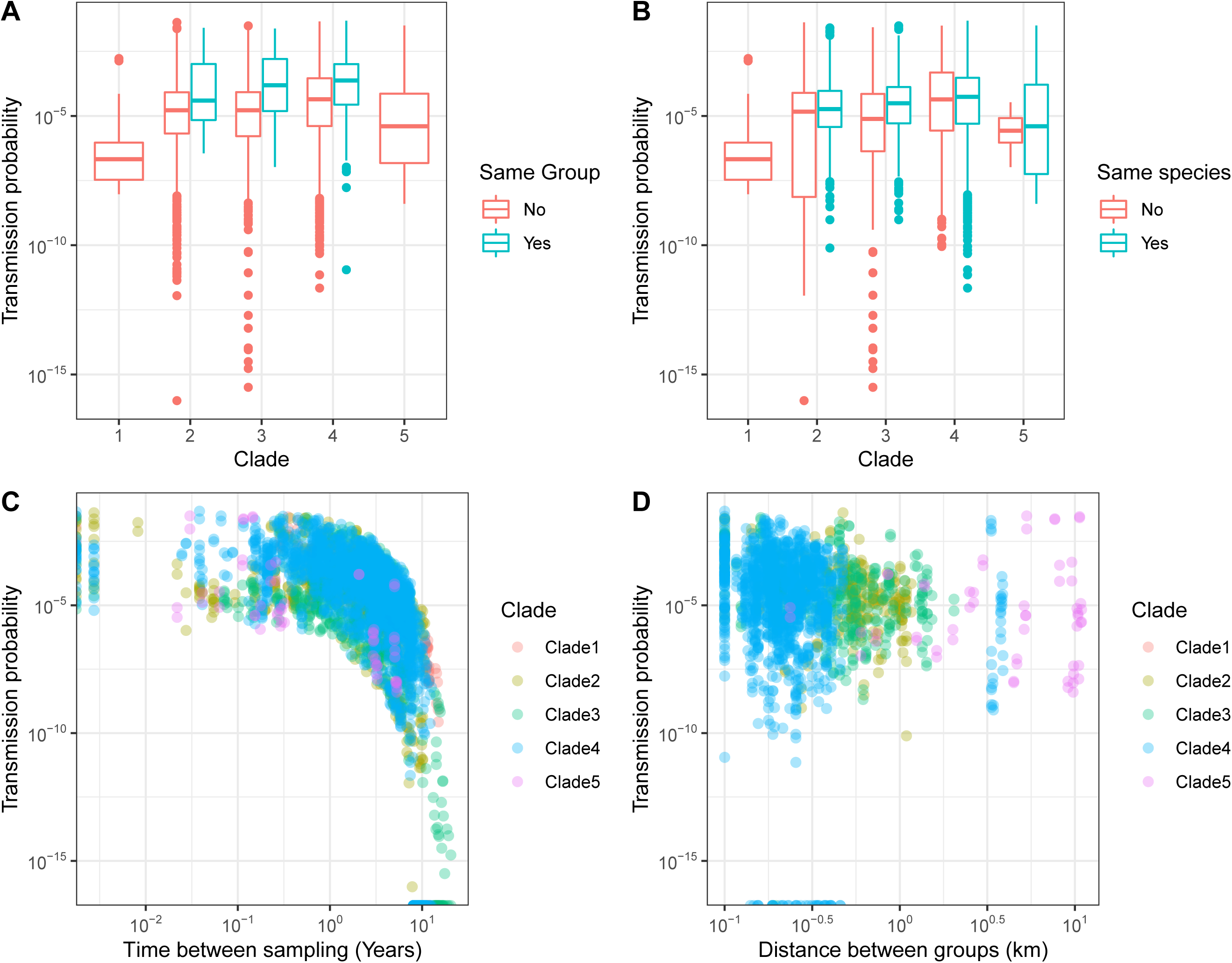
Epidemiological factors influence. Estimated transmission probability (y-axes) vs. same group category (panel A), same species (panel B), time between host pair sampling (panel C), and between-groups distance (panel D).

### 2.2 Transmission trees

In order to identify the most likely transmission trees, we assigned a weight (*W*_*AB*_) to each potential within-clade transmission between *A* and *B*, calculated by dividing the transmission probability P_A→B_ by the sum of all transmission probabilities in the same clade. Then, we started building the tree by drawing transmission pairs with probability depending on their weights, until the transmission tree was completed (i.e. only one host was left without a transmission source, representing the tree’s root). Assuming only a single source for each infection, we discarded all implausible transmissions resulting from previous selections (i.e. if a selected transmission was from *A* to *B*, then *B* to *A* was ruled out, as well as the transmission from other hosts to B), and also avoiding loops. For each clade we built 10,000 stochastic transmission trees, and for all trees, we calculated the *tree likelihood* (*L*) as follow:

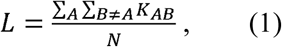

where *N* is the total number of transmissions in the tree, and

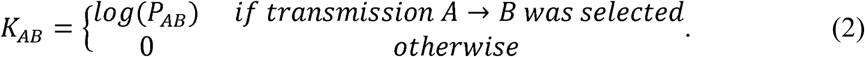

Here we report the results for clade 3 only. In this case, the median[range] of the stochastic trees likelihood was –7.85[–10.14, –6.91], and Fig 3 (left panel) shows the most likely transmission tree for this clade (i.e. corresponding to tree’s likelihood of –6.91).

**Fig 3.**
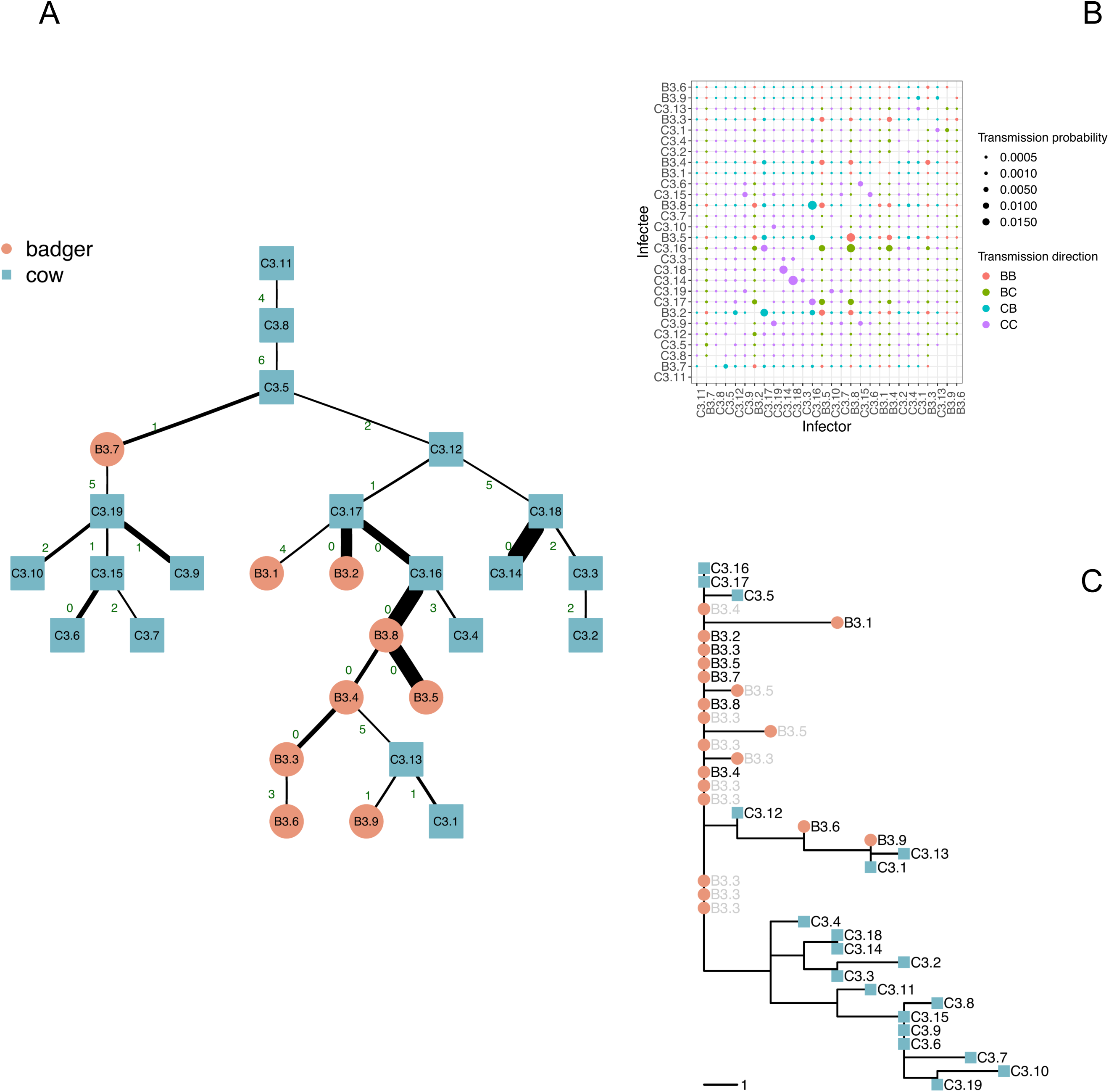
Probability matrix and transmission trees. A: transmission tree for clade 3 (top to bottom). Blue squares represent cattle, while red circles represent badgers. For each represented transmission *i* → *j*, the edge label indicates the total SNP distance between the isolates sampled in *i* and *j*, while line thickness represents the transmission probability *P*_*(i*→*j)*_. B: transmission probability for clade 3 pairs, the dot size is proportional to probability, and shade is proportional to the time elapsed between when the two sequences were sampled (black corresponds to the same day). C: clade 3 phylogenetic tree (grey-labelled strains were not used because multiple strains, identified by having the same label, were sampled from the same individual potentially at different times).

Transmission pairs associated with low SNP distance are, as expected, typically characterized by a higher probability (thick lines in Fig 3, left panel). However the inclusion of sampling time in the inference sometimes indicates that the individual with extra generated SNPs is more likely to be the source than the individual without them (see for example, C3.13 and C3.1, albeit with relatively low probability). As expected, where individuals have little SNP differentiation and similar sampling times, the transmission probabilities are similar and directionality is difficult to discern (e.g. a sub-cluster formed by individuals B3.2, B3.5, B3.8, C3.16, and C3.17; see Fig 3, upper-right panel). In contrast, where times intervals are long (e.g. individual C3.11 was sampled a decade earlier than others in its clade) the probability of transmitting *M. bovis* to other members of the clade was very low (ranging from 10^−16.1^ to 10^−7.8^), indicative of missing infected individuals. As shown in S1 text (Fig S3.1), the low and variable substitution rate for *M. bovis* mean that this can be true even for individuals with zero SNP distance but that were sampled more than four years apart, with the probability of an intermediary being higher than for direct transmission, and with the threshold decreased as the number of divergent SNPs increased. Three of the selected transmissions of the tree reported in Figure 3 (C3.11→B3.7, B3.8→C3.13, and C3.5→C3.3) had a difference in sampling times (Δt) that was higher than the threshold showed in S1 text (Fig S3.1), thus transmission was more likely to have been mediated by missing hosts than via a direct route.

### 2.3 Alternative bottle-neck model

In the model described above we assumed that SNP substitutions within an infected host can occur during latency (i.e. after infection but before infectiousness onset), so that the pathogen population bottleneck occurs at the point of infection, where only a very small number of bacteria are transmitted. It is however possible that the bacteria only replicate appreciably once active infection has started, and that in the latent stage, pathogen evolution and therefore substitution rates are low or zero. Although this issue has not been explored yet for *M. bovis*, controlled experiments with the closely related *M. tuberculosis* showed no variation in substitution rates in the latent and active disease stages in macaques ^35^; a further study in a population of infected humans found low growth and substitution rates during latency ^36^. While the former observation would imply a pathogen population bottleneck at the infection stage (i.e. when the host state changes to Exposed), the latter implies a bottleneck at the end of the latency stage (i.e. Infectious state) and thus no definitive model for the within-host evolution bottleneck exists.

Here, we explored the implications of different population bottleneck models by calculating the pairwise transmission likelihood with two alternative within-host models. Specifically, we allowed divergent SNPs to appear either from the point of infection, or at the infectious (I) stage only (i.e. setting the mutation rate for Exposed individuals to 0) for one of the clades (clade 3).

We showed that although the average transmission likelihood had not changed, and the lower estimates did not change much using the alternative model, for many of the more likely transmission pairs there was a substantially stronger effect (Fig 4). The tree reported in Fig 5 (panel A) shows that three selected but unlikely transmission pairs were consistent in both models (e.g. C3.11→C3.8, C3.12→C3.18, C3.17→B3.1). Differences in the likelihood of transmission events were also identified even when the strains were closely related (e.g. B3.1 was infected by C3.17 with the first model, and by C3.16 with the alternative one). These differences were due to the ability of this approach to discriminate between host pairs with similar SNPs distance (0 or 1). This was particularly evident for the triad formed by C3.16, B3.8 and B3.5, all at 0 SNPs distance between one another. While the original model inferred the transmission chain C3.16→B3.8→B3.5, the alternative model inferred that C3.16 infected both the B3.8 and B3.5. The triad formed by B3.9, C3.1 and C3.13 could be considered similarly, since the original model inferred C3.13→C3.1 and C3.13→B3.9, whilst the alternative one inferred B3.9→C3.1→C3.13. However, some clusters where SNP distances were close were robust to the bottleneck assumption, in particular those formed by C3.2, C3.3, C3.18, and C3.14, and by C3.6, C3.7, C3.9, C3.10, C3.15, and C.19, although in the latter the position of many hosts and the inferred direction of transmission differed. These clusters were also consistent with the phylogenetic tree shown in Fig 2, bottom-right panel.

**Figure 4.**
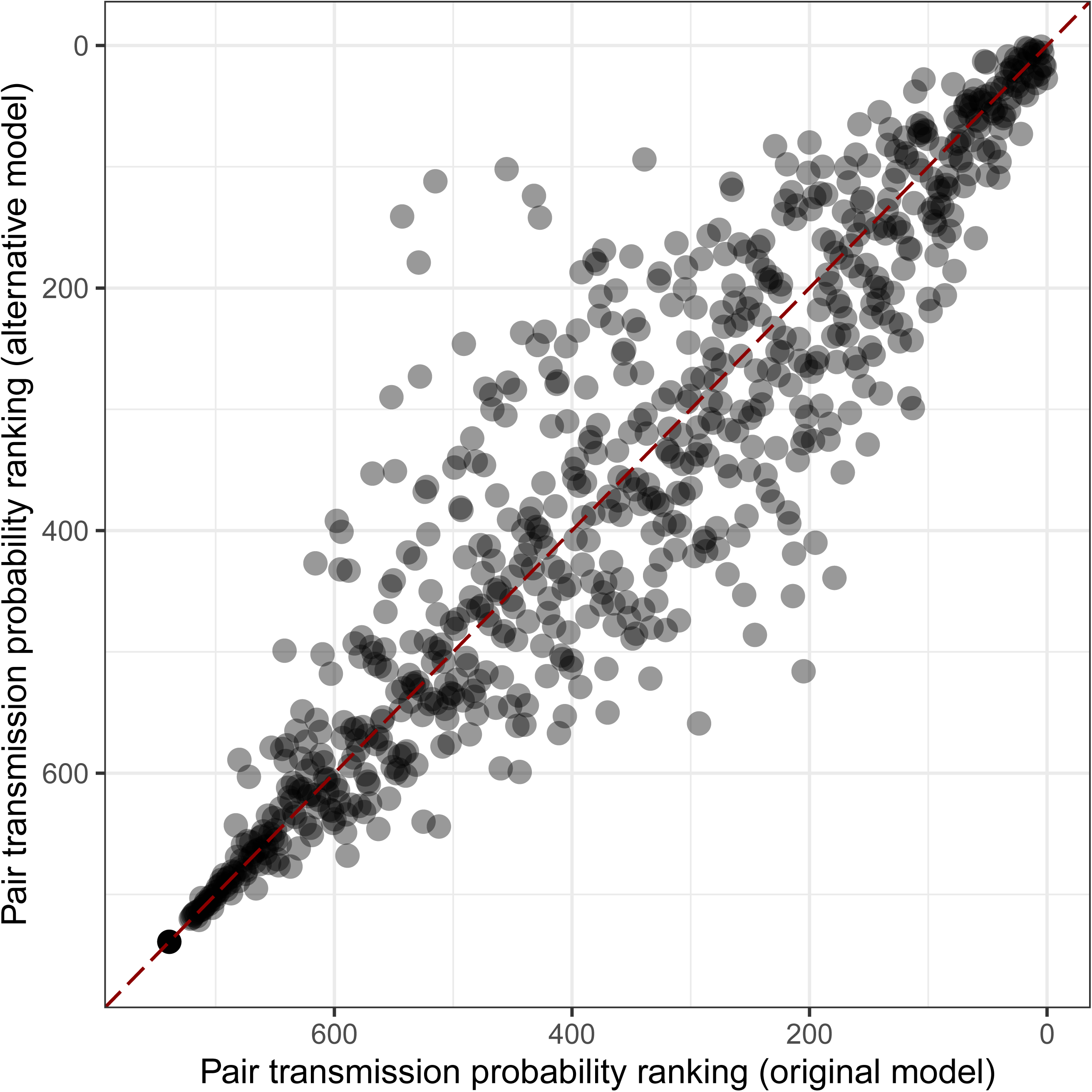
Transmission probabilities ranking with different bottleneck assumption. Comparison of the pairwise transmission probabilities ranking for each hosts pair using the original model (SNPs substitution allowed during the exposed stage, pathogen population bottleneck at infection) and an alternative within-host model (SNPs substitution allowed at infectious stage: pathogen population bottleneck after latent stage).

**Fig 5.**
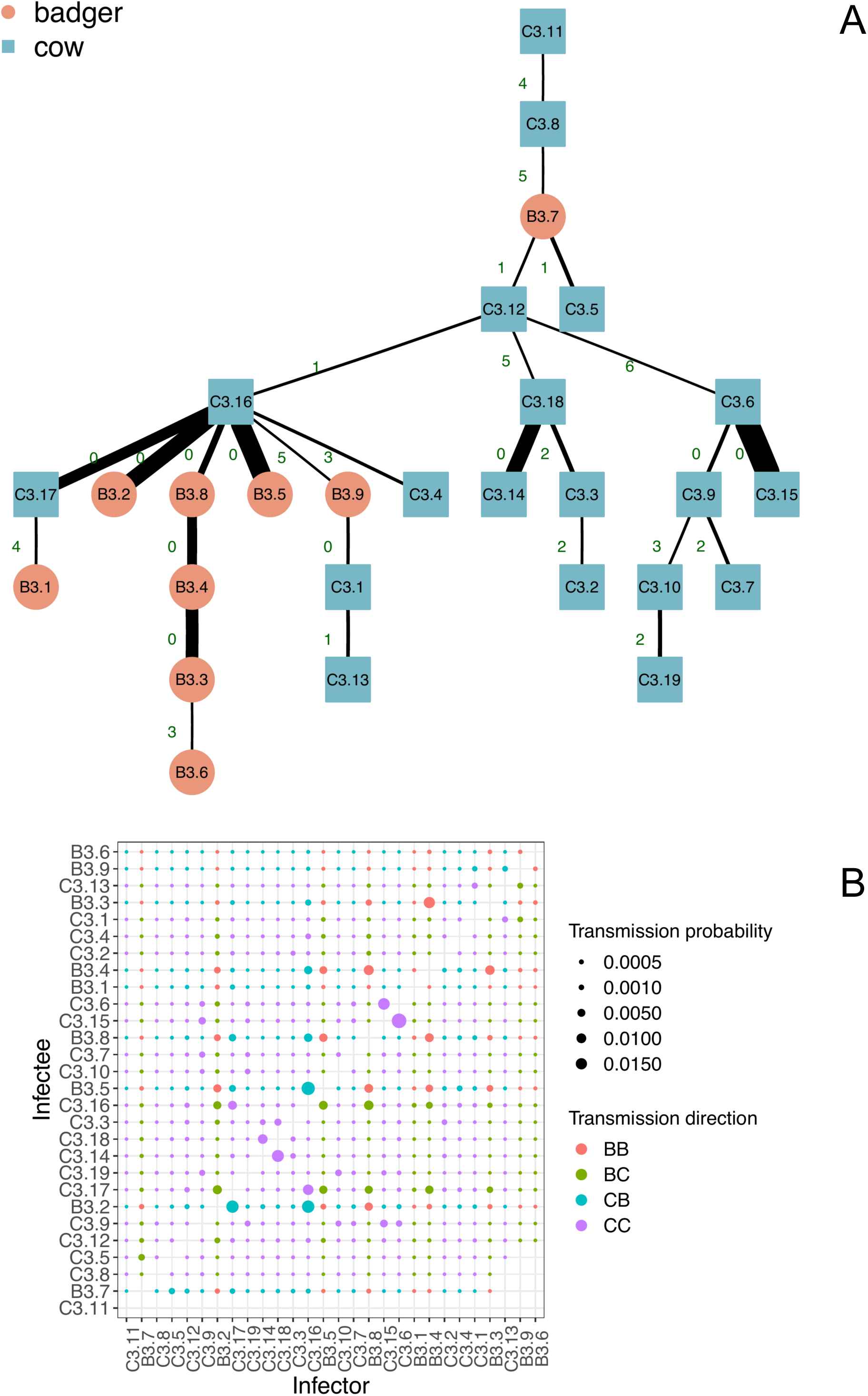
Transmission tree with alternative pathogen bottleneck model. Transmission tree (A, top to bottom) and transmission probability matrix (B) for clade 3, assuming an alternative model where SNPs substitution is allowed during the infectious period only (see Fig 3 caption for full description).

## 3. Discussion

In infectious disease epidemiology, the importance of considering processes at different scales for disentangling the dynamics of outbreaks is well recognised ^37,38^. Recent improvements in techniques to obtain pathogen genomic information have resulted in a number of advanced methods that can use genomic and epidemiological data to disentangle the roles of different contributing processes, including the infectious contact network ^15,25,39–41^. However, while many of these methods are suited to the analysis of rapidly evolving pathogens such as RNA viruses(i.e. HIV, SARs, Foot-and-Mouth Disease, Ebola), applying the same methods to slowly evolving and spreading pathogens is more challenging, due to the low signal-to-noise ratio (i.e. in SNPs) when comparing different strains ^26,37,42^. Chronic bacterial diseases usually present long and variable latency periods that increase uncertainty around the timing of transmission events, and also contribute to partially sampled outbreaks, as the infected host can be removed from the population by other causes before exhibiting the symptoms of infection ^43^.

Here, we have presented a model to study infectious diseases dynamics at a forensic level, combining within-host evolution of *M. bovis* and between-host transmission, which relies on genomic information and basic epidemiological data only. Since pathogen evolution and infection processes are intimately related, we showed that combining the sampling time (often the only epidemiological information available), whole genome sequence data for the bacteria, and a simple model for disease progression (here incorporating an exposed stage before infectiousness) can help discriminate amongst possible transmission pathways. This was possible even amongst groups of individuals infected with highly similar strains (i.e. fewer than five SNPs between them), even though it was not always possible to definitively identify who-infected-whom. As expected, a change in the within-host scale dynamics and therefore the pathogen population bottleneck can influence this result. Thus where information about within-host pathogen dynamics is available, these data should be used to inform the model. Where such data are not available, our method can be used to identify when inferred transmission pathways are robust to different bottleneck assumptions.

Our method exploits a crucial feature of the combined genetic and temporal data: while the time of sampling is useful to identify the probability boundaries for a possible transmission pair (broadly speaking, the farther apart in time for the sample dates, the less likely a transmission event), the genetic distance between the bacterial genomes embeds the combination of temporal distance and transmission distance over the network of potential contacts, parsed by the rate of pathogen within-host evolution. The inclusion of a latent stage further informs the inference (for example the inclusion of the latent stage reduces the probability of infectious contact if the sample dates and therefore the infection dates are too close together), and therefore an informed model should be better able to estimate inter-generation times and times of infection than approaches that do not consider the epidemiology.

Unlike many other approaches that consider the population as a whole, here we adopted a pairwise approach to estimation of parameters. A similar forensic approach was used by Campbell and colleagues in order to develop *outbreaker2*, which considered temporal, contact network, and genomic data to infer transmission events, although it did not consider the transmission process as we do here ^44^. While on the one hand pairwise approaches substantially simplify the computation, on the other they introduce dependency problems that might bias the transmission likelihood calculations ^44^. Our method is reliant on unbiased sampling, and it does not consider any higher order interactions that might be embedded in the system. Such interactions might be important if, for example, there are substantial interactions between the infection pressures from two individuals on a third. Counterbalancing this is the exactness of the KFE approach.

While earlier results show that in the Woodchester Park badger dataset geographical distance between *M. bovis* isolates is an important predictor of their genetic distance ^34^, here we show that this relationship is not a reliable indicator of the transmission likelihood. However, as we consider only pairwise interactions in the present analyses, it may be that the relationship with distance would be recovered in an analysis of longer chains. As expected, the social group provided a strong signal. These results suggest that, for this context, contact networks could be better informed by social interactions than by spatial distance. These results were obtained despite our methods suggesting many missing links in the obtained transmission trees, as we were able to detect the branch in the transmission trees where it is more likely that one or more infected hosts were not sampled. It is important to consider that the number of unsampled individuals is not known, and so it is always possible that an unsampled individual was directly involved. However, the density of sampling of the Woodchester Park badger population was high, with animals being trapped and tested for *M. bovis* infection on average twice every year ^45^. Even considering the relatively low sensitivity of the testing approach ^46^ we would still expect that our highly likely pairs would have more closely related infections compared to our less likely pairs. That we were able to identify circumstances when an intermediate host was likely is particularly notable given the low level of diversity in *M. bovis* genomic data, and the fact that we used only up to five SNPs in our estimates. The analyses presented here follow a simplistic representation of transmission dynamics, but could easily be expanded within the KFE framework, albeit at increased computational cost. For example, we did not account for events other than the sequence sampling. Since UK cattle are regularly tested for bTB as part of the routine national control program, we could include the probability of being infected but not detected on the date of a negative test prior to a positive one. Other epidemiological data (distance, population groups, species, etc.) could also be explicitly embedded, and might identify, for example, differences in estimated parameter posteriors for different species combinations.

In our calculations, we considered values of SNP difference up to five. The choice of this low SNP threshold was mainly for computational reasons, as for every extra SNP the size of the matrix used to solve the KFEs increases exponentially, thus slowing the calculation, as illustrated in equation (1). However, Walker and co-authors ^47^ showed that for human TB five SNPs is likely sufficient to discriminate between likely and unlikely transmission events. Our results showed a similar pattern with the probability dropping below 10^−5^ past the fourth SNP (Fig 1). In order to build the transmission tree we used a simple but intuitive algorithm compared to other methods in the literature (^25^, and references therein), which allowed us to identify the poorly supported transmissions where unsampled infected hosts were more likely to be involved. A more sophisticated algorithm for building the transmission trees would provide additional insight but at computational costs.

The objective of this study was to disentangle the interactions between processes happening at the within-host scale and at the population level, in order to refine the representation of the potential transmission network, compared to considering SNP distance alone when limited metadata and only a few samples are available. Using the KFEs in a novel analytical approach, we have shown how accounting for the evolution of *M. bovis* strains and incorporating an epidemiological model can successfully distinguish between many likely and unlikely transmission scenarios, despite the low genetic variability observed due to the slow and variable substitution rate that characterizes *M. bovis*, and with only a limited number of samples. Our KFE method is flexible and precise, and could be applied to other chronic infections where identifying who infected whom can be difficult, such as Johne’s disease (*Mycobacterium avium paratuberculosis* infection) in cattle, or leprosy in humans, contributing to an improved understanding of the role of within-host evolution and epidemiological dynamics in inferring contact patterns.

## 4. Methods

### 4.1. The Pairwise Kolmogorov Forward Equations (KFEs) with within-host dynamics

We specify a three-state Susceptible-Exposed-Infectious (SEI) model as being appropriate to the epidemiology of *M. bovis* infection dynamics ^43,48–53^. Here, *Susceptible* individuals become *Exposed* (but not infectious) after a successful transmission from an *Infectious* individual, and then move to an *Infectious* state after the latency period. The transmission rate and the E to I progression rate are represented by β and σ, respectively (thus 1/σ is the average duration of the E stage). We consider within-host evolution in parallel with disease progression in the dynamic model, in order to account for differences between the observed and transmitted lineages of bacteria in a host ^10^ and to explore the potential impact of variation in population dynamics at the within-host level. For simplicity we consider all nucleotide transitions to be equally likely (i.e. rate of A→T is the same as rate of C→G, etc.).

Strain evolution is modelled as a dynamic process occurring simultaneously alongside the infection progression, with the number of SNPs indicated by the superscript *k* (thus the full host state is denoted as 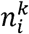, with *n*_*i*_ representing the epidemiological status) and these SNPs generated at a fixed *substitution rate* (μ). Here, we refer to the strain harboured by a host before it is transmitted to another as *ancestral*, and it is denoted by *k = 0* (i.e.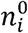). When *k*>*0* it denotes the number of SNPs 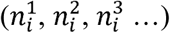 in the mutant strains. Following previous researchers, we have labelled the extra SNPs in a given sampled mutant strain as *divergent SNPs* ^26^. These SNPs does not include the substitutions occurring before the transmission, as we consider these common to both hosts strains. In the case of a pair of hosts, the KFE system is defined as follows:

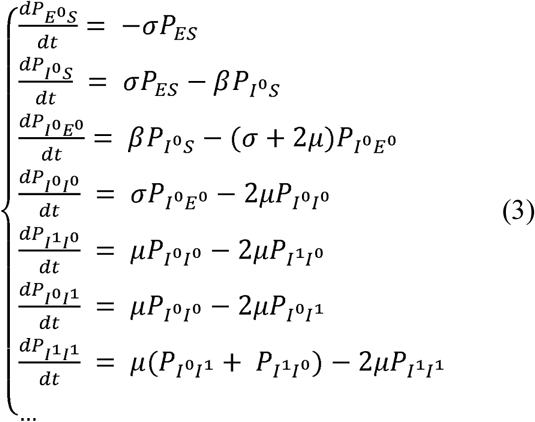

Here, 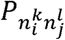 is the probability of *A* being in status *n*_*i*_ and infectious and *B* being in status *n*_*j*_, with *k* and *l* divergent SNPs, respectively. For a full derivation of the KFEs see the Supplementary material S1.

Thanks to this system, we can calculate the exact probability for two hosts in any possible combination of infection states and for any number of divergent SNPs. Hence, assuming that host *A* was first exposed to the infection at time *t*_*0*_, host *B* was infected by the former, and pathogen strains from both hosts were sampled at time *t*_*T*_ (= *t*_*A.*_= *t*_*B*_),we can numerically solve the system (1) from *t*_*0*_ and *t*_*T*_ to obtain the transmission probability:

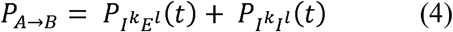

with *t = t*_*T*_ – *t*_*0*_.

In case the sampling time differs (*t*_*A.*_≠ *t*_*B*_), we have to calculate the transmission probability in two steps. Assuming *t*_*A*_<*t*_*B*_, we can solve the system (1) from time *t*_*0*_ until A was removed at *t*_*A*_ (removal/sampling time). Then, we could plug the results into a single-host KFE model and solve this one between *t*_*A*_ and *t*_*B*_ (for the one-host KFE system see Supplementary material S1). An analogous reasoning holds for the case *t*_*B*_<*t*_*A*_.

The equations in (1) are linear, and so in principle it is trivial to find the solution of the P_*A*→*B*_.However, given the complexity of the system and number of equations increasing with the number of SNPs considered, we opted for a numerical solution implemented in R^54^, with package *deSolve*^55^.

### 4.2. Application of the pairwise KFE model

Clinical samples are taken from each captured badger and undergo microbiological culture to attempt to isolate *M. bovis*. Whole genome sequences from isolates collected since 2000 were made available for the present study. In addition, *M. bovis* sequences sourced from cattle on neighbouring farms were also made available. These farms were sampled as part of the routine bTB control and eradication program, and the associated testing information is stored in the APHA cattle testing (SAM) database ^56^. Following the analysis by Crispell et al.^34^, we maintained the Woodchester Park infected population divided into five main clades (see Table 1). The clades were defined as containing highly similar *M. bovis* sequences (all isolates within 10 SNPs of one another) sourced from cattle and badgers. We used this information as a proxy for evidence of a high likelihood of a contact. We conducted our analysis on each clade separately, except for clade 4 which was subdivided into a further four subclades for computational reasons. For badgers yielding more than one *M. bovis* sequence, we only considered the most recent one. We then counted the differences between each pair of strains, to obtain the number of divergent SNPs. This process was applied to all pairs of individuals present in each clade (or sub-clade). For the sake of simplicity, we labelled all individuals according to their species (B for badgers and C for cattle), main clade (1 to 5), and a sequential integer number.

**Table 1.** Woodchester Park (UK) *M. bovis* sequences and clades (based on Crispell et al.^34^).

| Clade | Sub-clade | # *M. bovis* sequences | # *M. bovis* sequences from badgers | # *M. bovis* sequences from cattle | # hosts | # badgers | # cattle |
| --- | --- | --- | --- | --- | --- | --- | --- |
| Clade 1 | - | 11 | 4 | 7 | 8 | 1 | 7 |
| Clade 2 | - | 32 | 10 | 22 | 27 | 5 | 22 |
| Clade 3 | - | 39 | 20 | 19 | 28 | 9 | 19 |
| Clade 4 | (total) | 170 | 156 | 14 | 107 | 93 | 14 |
| Clade 4 | a | 27 | 24 | 3 | 16 | 13 | 3 |
| Clade 4 | b | 29 | 27 | 2 | 18 | 16 | 2 |
| Clade 4 | c | 24 | 23 | 1 | 15 | 14 | 1 |
| Clade 4 | d | 28 | 26 | 2 | 22 | 20 | 2 |
| Clade 4 | e | 10 | 8 | 2 | 6 | 4 | 2 |
| Clade 4 | f | 19 | 17 | 2 | 11 | 9 | 2 |
| Clade 4 | g | 33 | 31 | 2 | 19 | 17 | 2 |
| Clade 5 | - | 8 | 1 | 7 | 8 | 1 | 7 |

For each pair of sampled individuals in each clade (or sub-clade), we independently calculated the probability that one host infected the other (considering both directions), given the timings of sampling for the two hosts, and the observed divergent SNPs at the respective time of sampling. We defined a constant death rate, based on the observation that less than 0.1% of individuals would survive past 8-years of age (*maximum lifespan*) for both cattle ^57^ and badgers ^58^.

As the epidemiological parameters for bTB are highly uncertain, instead of using a single parameter value we drew them from distributions informed by the recent literature. We then allowed variability in the probability calculation by moving each parameter according to a Gaussian kernel (mean the previous iteration values, sd = 5%) for 1,000 iterations, each time calculating the pairwise transmission probability. We finally selected the combination of parameters which resulted in the highest transmission probability.

The distribution for the substitution rate (*μ*) was a beta-PERT, with mode, minimum and maximum respectively set to 0.31, 0.1 and 0.94 base pair × genome × year. These values corresponded to the average, minimum, and maximum of literature estimates from other systems ^27,59–61^. We assigned a similar prior to the latency period (*1/σ*), based on the published literature for data relevant to the geographical area (south-east England). In this case we used a beta-PERT distribution, with mode, minimum and maximum respectively set to 348, 116 and 827.5 days ^51,52^. Finally, we used a uniform distribution for the contact rate: *β* ∈ *U*(0, 0.1) × contact × year. This was set to include the transmission rate estimates in ^51,52^, but widened to account for the fact that the pairwise infection rate in our model assumes that contact exists (i.e. the population level parameters consider both probability of contact, and infection rate given that contact).

### Data availability

All WGS data used for these analyses have been uploaded to the National Centre for Biotechnology Information Short Read Archive (NCBI-SRA: PRJNA523164). Because of the sensitivity of the associated metadata, only the sampling date and species will be provided with these sequences.

## Acknowledgements

This work was supported by BBSRC (grants BB/P010598/1, BB/L010569/1, and BB/L010569/2), SFI (grant 16/BBSRC/3390), and DEFRA.

## Authors contributions

GR contributed to the model development, conducted the analyses, and wrote the manuscript. JC advised on the analysis, and contributed to the data generation. DB contributed to the data generation. SJL advised on the analysis. RJD helped to conceive the study and help to interpretation of the data. RRK conceived the study, contributed to the model development and advised on the analysis. All authors advised on the manuscript.

## Supplementary material

Complementary analyses including the full derivation of the pairwise Kolmogorov Forward Equations (S1), the analysis of simulated data (S2), and the pairwise vs. triplet Kolmogorov Forward Equations models (S3).

