## Supplementary material for "Identifying likely transmission pairs with pathogen sequence data using Kolmogorov Forward Equations; an application to *M.bovis* in cattle and badgers"

#### S1. Kolmogorov Forward Equation derivation

##### *(a) Within-host dynamics*

The proposed method uses the KFEs to represent the dynamics of an epidemiological system[1], but here addressing the additional complexity of embedded pathogen evolution. Instead of recording only the typical number (or fraction) of individuals in a given infection state (e.g. number of Susceptible, Infectious and Recovered in the SIR model case) as is the case for the usual compartmental model formulation of transmission dynamics, the KFEs describe the probability of the system having a given status, with an exact number of individuals in each infected state. If we consider a population of 50 individuals and an SIR model, the KFEs describe the probabilities: $P_{(S = 49, I = 1, R = 0)}$, $P_{(S = 48, I = 2, R = 0)}$, $P_{(S = 48, I = 1, R = 1)}$, etc. Keeling and Ross[2] showed the KFEs can be used to study the behaviour of pathogens in small and well-mixed populations. In the present analysis we followed Sharkey[3], who used this method to describe infection dynamics starting at the individual level.

By generalizing the epidemiological compartmental models, given an individual host and an infectious disease characterized by $N$ number of possible infected states, we can write a set of equations describing the probability the host transitions from state $n_0$ to state $n_i$ with $i \in (1, N)$ as:

$$\begin{cases} \frac{dP_{n_0}}{dt} = \dot{P}_S = -\varepsilon P_S \\ \frac{dP_{n_i}}{dt} = \sigma_{i-1}P_{i-1} - \sigma_i P_i \end{cases} \quad . \qquad (1)$$

Here, ε represents an external *force of infection* (which depend on the number of other infected host in the population of interest), while $\sigma_i$ represents the *transition rate* from status $n_i$ to $n_{i+1}$.

During pathogen replication, errors in DNA copying (or RNA copying if RNA genomes) result in nucleotide substitutions, insertions or deletions. Generally these single nucleotide polymorphisms (SNPs) are considered to be neutral in bacterial species within epidemics[4]. For our model here, we will assume that SNPs are neutral and do not consider the cases where a mutation could significantly enhance the pathogen fitness within a particular environment (e.g. we are not considering SNPs which confer drug-resistance, or special adaptation to a new host species).

Whole genome sequencing (WGS) can be used to detect polymorphisms, and the presence or absence of common or divergent substitutions in a pair of mutant strains can be used to estimate their relatedness. Assuming to know the strain an individual is infected with, we can include strain evolution as a dynamic process occurring simultaneously alongside the infection progression, with the number of SNPs indicated by the superscript *k* (thus the full host state is denoted as $n_i^k$). Here, we referred to original strain circulating in a host and before it is transmitted to another as *ancestral*, and it is denoted by $k = 0$ (i.e. $n_i^0$). When $k>0$ it denotes the number of SNPs ($n_i^1$, $n_i^2$, $n_i^3$ …) in the mutant strains. We defined the extra SNPs in a given sampled mutant strain as *divergent SNPs* (as they make the mutant strain diverge *ancestral strain*). Although this differentiation is not particularly useful in the single-individual framework, it will become as we move to consider a pair of hosts (see Figure S1.1, left). In the case of the single-host described general model, we can extend equation (1) by adding the within-host strain evolution to the disease progression as follows:

$$
\begin{cases}
\frac{dP_S}{dt} = -\varepsilon P_S \\
\frac{dP_{n_1^0}}{dt} = \varepsilon P_S - (\sigma_1 + \mu) P_{n_1^0} \\
\frac{dP_{n_1^1}}{dt} = \mu P_{n_1^0} - (\sigma_1 + \mu) P_{n_1^1} \\
\frac{dP_{n_i^0}}{dt} = \sigma_{i-1} P_{n_{i-1}^0} - (\sigma_i + \mu) P_{n_i^0} \\
\frac{dP_{n_i^k}}{dt} = \mu P_{n_i^{k-1}} + \sigma_i P_{n_{i-1}^k} - (\sigma_i + \mu) P_{n_i^k} \\
\dots
\end{cases}, \qquad (2)
$$

where μ corresponds to the *substitution rate*. This system implicitly implies three assumptions: (*i*) there is only one strain infecting a single host, (ii) within-host substitutions happens instantaneously and the time-to-event is exponentially distributed, and (iii) the effective population size is arbitrarily small.

For simplicity we do not differentiate between mutations to different nucleotides and assume that all nucleotide transitions are equally likely (i.e. rate of A → T is the same as rate of C → G, etc.). Assuming a host has been exposed to the infection at time $t^0$ and sampled at time $t^T$, we can calculate the probability of the strain showing any given number of SNPs by numerically solving equation (2) in this time interval (from $t^0$ to $t^T$).

#### *(b) SEI model and the pairwise KFEs*

For the purpose of this study, we described the equations using a three-state Susceptible-Exposed-Infectious (SEI) model (see main text, Materials and Methods section).

Considering this model implies assuming a maximum of $N = 2$ infectious states in which $n_o = S$, $n_1 = E$, $n_2 = I$, $\sigma_1 = \sigma$ (i.e. transition rate from E to I), $\sigma_2 = 0$. The equations for a single host then become (see Figure S1.1, right):

$$
\begin{cases}
\frac{dP_S}{dt} = -\varepsilon P_S \\
\frac{dP_{E^0}}{dt} = \varepsilon P_S - (\sigma + \mu)P_{E^0} \\
\frac{dP_{E^1}}{dt} = \mu P_{E^0} - (\sigma + \mu)P_{E^1} \\
\frac{dP_{I^0}}{dt} = \sigma P_{E^0} - (\sigma + \mu)P_{I^0} \\
\frac{dP_{I^1}}{dt} = \mu P_{I^0} + \sigma P_{E^0} - (\sigma + \mu)P_{I^1} \\
\dots
\end{cases} \quad . \qquad (3)
$$

After defining the equations for a single host, we can extend them for a host pair. In this case, we assume that, at the beginning of the calculation, one host is infected but not yet infectious (labelled *A*), while the second host is still susceptible (labelled *B*). A second necessary assumption is that individuals *A* and *B* can be in contact, and that *B* can only be infected by *A*. Thus, we removed the external forced of the infection and the transition from *S* to *E* for individual *B* is controlled by the infection rate $\beta$.

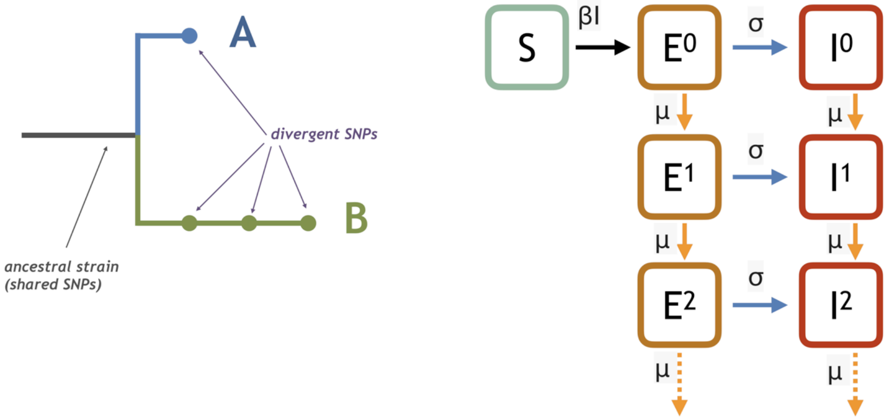


***Figure S1.1. Model schemes.*** *Left: the phylogenetic tree of two mutant strains respectively sampled in hosts A and B. The ancestral strain correspond and genome at the moment of the transmission (A→B or B→A). In this example, host A has 1 divergent SNP with respect to B, while host B has 3 with respect to A, therefore total SNP distance is 4. Right: the compartmental Susceptible-Exposed-Infectious model (SEI; $\beta$ represents the infection rate, $\sigma$ the transition rate from exposed to infectious, and $\mu$ the SNPs substitution rate). Since we are tracking the divergent SNPs only, we started counting them after the infection.*

Ignoring the strain evolution, we can write the equations for a host pair as follows:

$$
\begin{cases}
\frac{dP_{ES}}{dt} = -\sigma P_{ES} \\
\frac{dP_{IS}}{dt} = \sigma P_{ES} - \beta P_{IS} \\
\frac{dP_{IE}}{dt} = \beta P_{IS} - \sigma P_{IE} \\
\frac{dP_{II}}{dt} = \sigma P_{IE}
\end{cases}, \quad (4)
$$

where $P_{ij}$ is the probability of $A$ being in status $i$ and infectious and $B$ being in status $j$ (e.g. $P_{ES}$ correspond to $P_{ij}$ with A in status $i = E$ and B in status $j = S$).

The underlying assumption is that one host harboured a single pathogen strain at the time of infection, but further pathogen replication resulted in the generation of pathogen diversity. One lineage is then sampled in the 'source' host (or 'infector'), while the strain taken from the 'recipient' (or 'infectee') host comes from a different lineage and is likely to display additional diversity that distinguishes it from the first. Since eventual mutation happened in the ancestral strain prior to its transmission to the infectee did not contributed to the genetic distance between the two, this model does not consider them and it accounts only for the ones happened after the presumed infection.

By adding the strain evolution dynamics, we obtain the following system:

$$
\begin{cases}
\frac{dP_{E^0S}}{dt} = -\sigma P_{ES} \\
\frac{dP_{I^0S}}{dt} = \sigma P_{ES} - \beta P_{I^0S} \\
\frac{dP_{I^0E^0}}{dt} = \beta P_{I^0S} - (\sigma + 2\mu) P_{I^0E^0} \\
\frac{dP_{I^0I^0}}{dt} = \sigma P_{I^0E^0} - 2\mu P_{I^0I^0} \\
\frac{dP_{I^1I^0}}{dt} = \mu P_{I^0I^0} - 2\mu P_{I^1I^0} \\
\frac{dP_{I^0I^1}}{dt} = \mu P_{I^0I^0} - 2\mu P_{I^0I^1} \\
\frac{dP_{I^1I^1}}{dt} = \mu (P_{I^0I^1} + P_{I^1I^0}) - 2\mu P_{I^1I^1} \\
\ldots
\end{cases}. \quad (5)
$$

***(c) Transmission probability calculation***

The main goal is to use the KFEs to calculate the transmission probability $P_{A \to B}$ from an infector A to an infectee B. This translates into using the KFE system (5) to calculate:

$$P_{A \to B} = P_{I^k E^l}(t) + P_{I^k I^l}(t) \qquad (6)$$

with $t = t_T - t_0$, which corresponds to the time span between A infection ($t_0$) and both host sample ($t_T$ , i.e. the time of pathogen strains observation), with $k$ and $l$ representing the divergent SNPs of A and B strains, respectively.

At the initial time ($t_0$) we set the probability of the infector and infectee being, respectively, at the exposed (E) and susceptible (S) stage to one: $P_{E^0 S}(t_0) = 1$, while all the other system states probability is zero ($P_{I^0 S}(t_0) = P_{I^0 E^0}(t_0) = P_{I^0 I^0}(t_0) = P_{I^1 E^0}(t_0) = \cdots = 0$). From this point, it is straightforward to calculate the probability for any possible combination of divergent SNPs in the two hosts strains, and for any time forward.

In case the sampling times differs ($t_A \neq t_B$), we have to calculate $P_{A \to B}$ in two steps. If $t_A < t_B$, we can first use the system (5) from time $t_0$ until A was removed at $t_A$. Then, we have to sum the probabilities calculated with the former for any potential state of B as follows:

$$\begin{cases} P_{E^0}(t_A) = \sum_Z P_{ZE^0}(t_A) \\ P_{I^0}(t_A) = \sum_Z P_{ZI^0}(t_A) \\ P_{E^1}(t_A) = \sum_Z P_{ZE^1}(t_A) \\ \qquad \cdots \end{cases} , \qquad (7)$$

with Z representing a vector including all possible states of host A. Therefore, we can set the initial condition for a single-host KFE model like the one represented in (3), and solve this from $t_A$ to $t_B$. An analogous reasoning holds for the case $t_B < t_A$.

### S2. Simulated tree analysis

#### *(a) Transmission trees simulation*

In order to validate our methodology, we applied it to a simulated dataset. We simulated 50 transmission trees using the R package epinet[5] (model homogeneous Susceptible-Exposed-Infectious-Recovered, total population: 60 individual, $\beta = 0.00075$, exposed period: 8 months, infectious period 10 months, SNPs substitution rate 0.00001, maximum number of loci 1000). We assumed that the state Removed transition corresponded to the sampling. Once transmission trees were created, we imported them in ΠBuss[6] in order to generate a phylogenetic tree, with genetic distance measured in SNPs (saved in a *.fasta* file).
Among the 50 trees, we choose two trees with 10 or less infectious individuals for computational reasons, but showing genetic diversity (max SNPs distance ≥3). The two trees are reported in Figure S2.1.

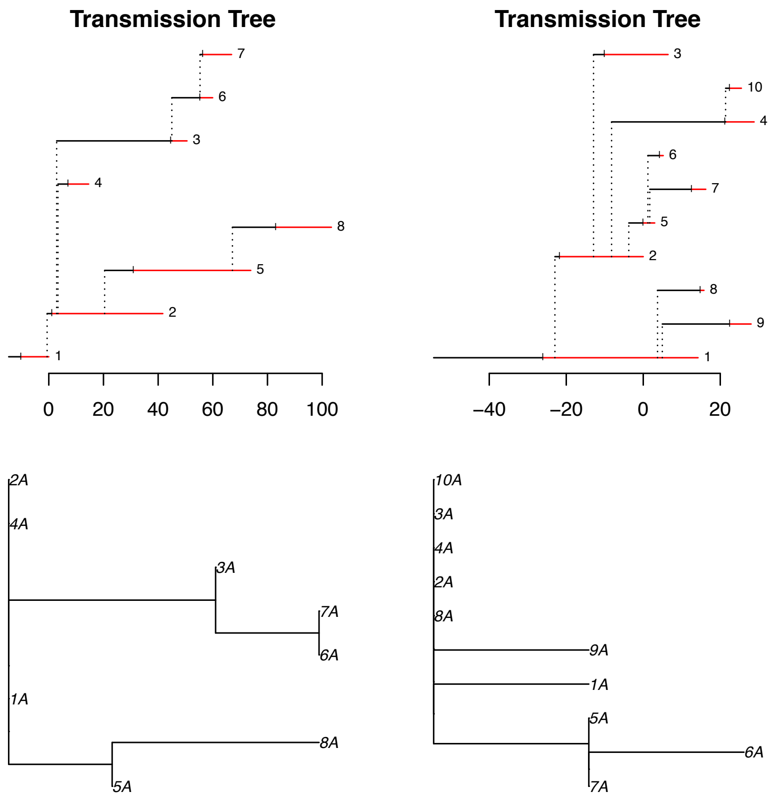


***Figure S2.1. Tree generated with epinet and ΠBuss.*** *Generated transmission trees (upper row) and corresponding phylogenetic trees (lower row).*

#### *(b) KFE analysis*

Following the methodology described in the main text Method and in S1, we analysed all transmission pairs with the Kolmogorov Forward Equations. In order to simulate parameters uncertainty, we used the same parameters space algorithm and a parameter distribution for each one (β: *betaPERT(0.00025, 0.00075, 0.0015)*, exposed period: *U(1, 24)*, SNPs substitution rate *U(0.000001, 0.001)*).

As reported in Figure S2.2 (panel A), for both trees the pairs SNP distance affects the calculated transmission probability. However, by plotting the probability against the difference in sampling time (Figure S2.1, panel B), we could observe that for similar strain (SNP distance = 0) the probability is higher for host pairs sampled closer in time, while as we increase the SNP distance, in most case the probability had a different behaviour with some increase for hosts sampled apart.

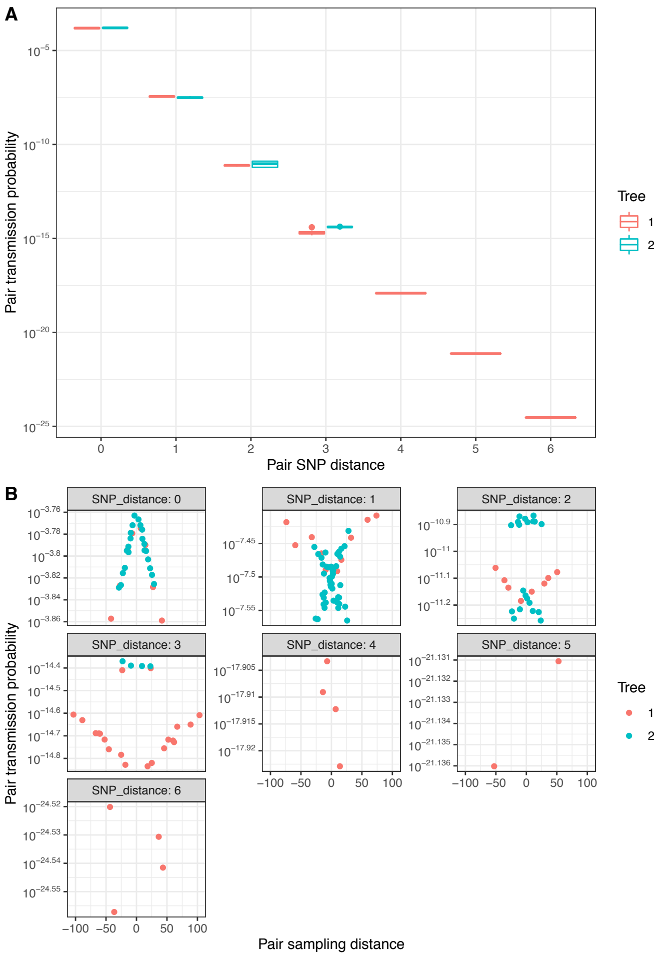


***Figure S2.2. Transmission probability calculated with the KFEs.*** *A: pairs transmission probability (y-axes) vs. SNP distance. B: pairs transmission probability (y-axes) vs. distance in time of sampling, with each panel representing a different SNP distance.*

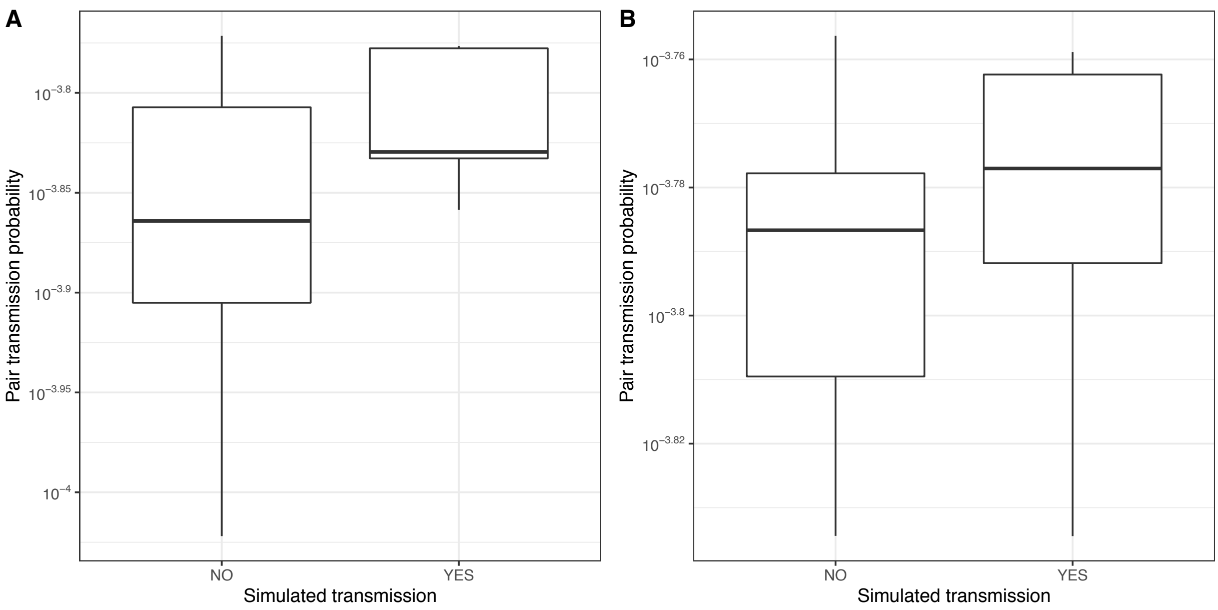


***Figure S2.3. Transmission probability for simulated transmission.*** *Comparison of transmission probability (y-axis) for pairs in the simulated tree 1 (panel A) and tree 2 (panel B). For each panel, the two boxplots represent the values for the simulated transmission vs the non-simulated ones.*

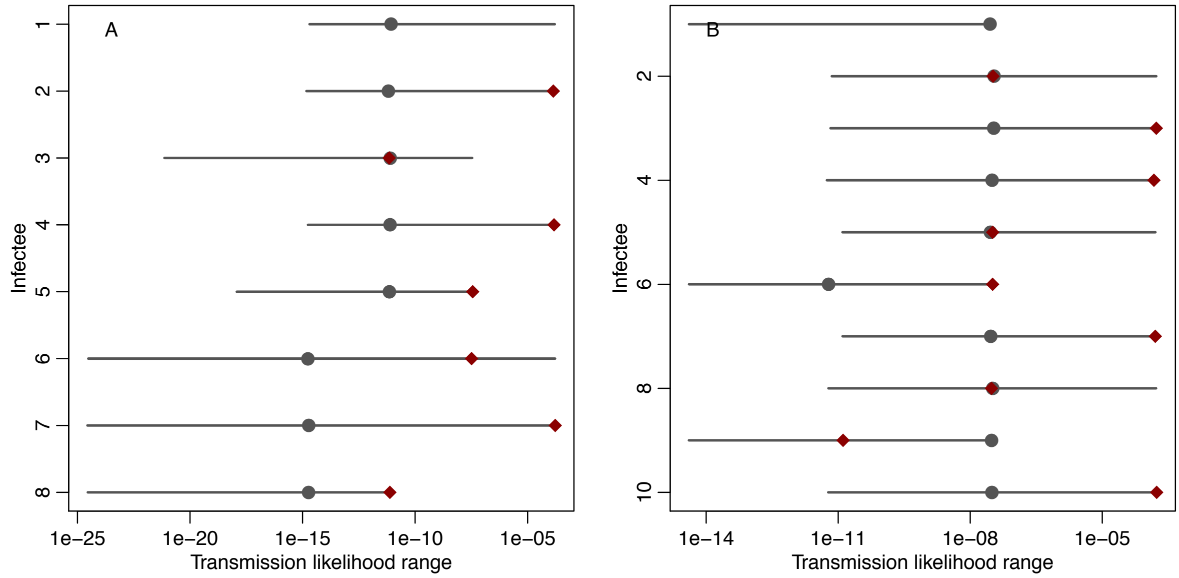


***Figure S2.4. Transmission probability for each infectee.*** *For each individual in the transmission tree, we compared the probabilities when it is on the receiving end of the transmission (i.e. when it is the infectee). The grey line represents the transmission probability range, grey dots the median, and red diamond the value for the simulated transmission (for both trees the root is individual 1, therefore it does not have a simulated transmission value).*

When comparing the simulated transmissions with the non-simulated ones, we observed higher probability values in the former for both trees (Figure S2.3). Moreover, when comparing the transmission probabilities for each infectee in isolation (Figure S2.4), in most cases the simulated one was the most likely transmission to the same host (five out of seven for Tree 1, panel A, and five out of nine in Tree 2, panel B), and only in one case was the simulated transmission substantially less likely the median of the other ones (individual 9, Tree2).

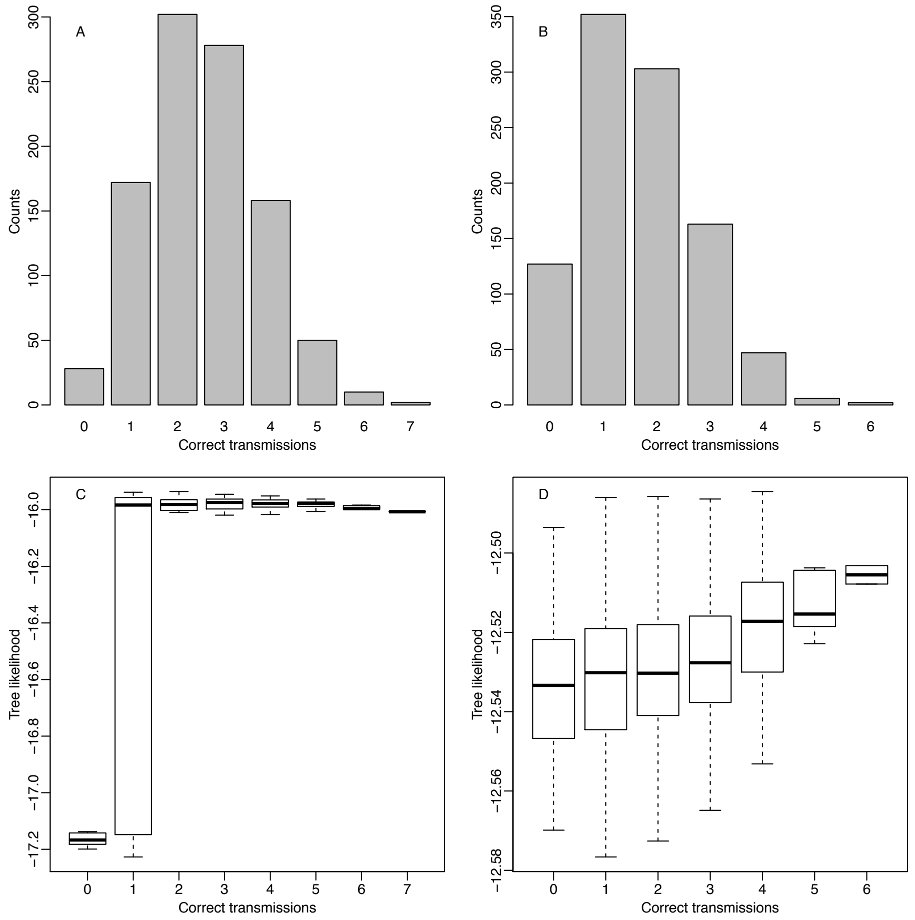


***Figure S2.5. Correct transmissions in the stochastically selected transmission trees.*** *For each of the 1,000 stochastically selected transmission trees, we calculated the number of simulated transmission in Tree 1 and Tree 2 (panel A and B, respectively). Lower rows shows the trees likelihood distribution for selected trees depending on the number of correct simulated transmissions (panel C and D for Tree 1 and 2, respectively).*

In order to check the transmission tree reconstruction procedure (main text, section 2.2) we run 1,000 stochastic reconstruction of the simulated trees. As showed in Figure S2.5, for Tree 1 (panels A and C) we were able to find the correct configuration (seven out of seven correct transmissions). As the number of correct transmission increases, the median likelihood increases as well, until five out of seven correct transmissions were selected. Past that point, the average likelihood decreases, probably because of the lower number of configurations. For Tree 2 the likelihood increases until there are six out seven correct transmission pairs, but out of the 1,000 stochastically reconstructed trees, we did not observe any with seven, eight or nine correctly identified transmission pairs. This was due to the lower score of the correct transmission to infectees 2, 6, and 9 (see Figure S2.4).

Overall, we can conclude that the KFEs methodology provide good results in providing hidden information by combining the genetic distance and epidemiological information. Tree reconstruction could be improved, although as the number of similar strains increases, it becomes more difficult to identity the correct transmissions.

**Section 3. Pairwise vs. triplet KFE**

Using Kolmogorov Forward Equations (KFEs) we first considered the likelihood of transmission between a pair of hosts $A$ and $B$, and subsequently examined the likelihood that a third unsampled individual ($U$) was also involved, by calculating the probability of a transmission between $A$ and $B$ through $U$ (Figure S3.1). By comparing the estimated transmission probability of the three-individual chain ($P_{A \to U \to B}$) with the pair chain ($P_{A \to B}$) we determined whether an intermediate was likely to have been missed in the transmission chain.

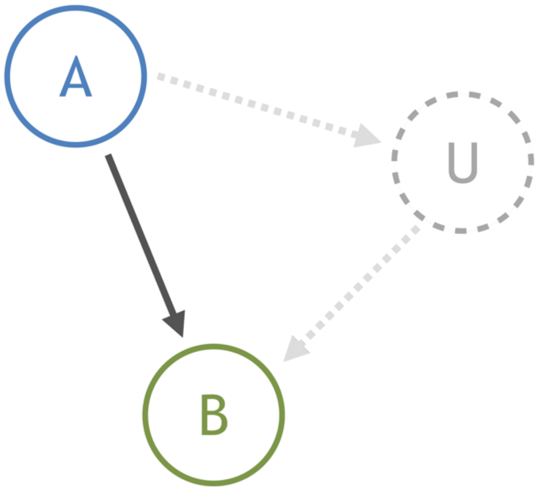


*Figure S3.1. The triad problem: given two observed infected hosts (A and B), how likely is it that transmission was direct or via a third unsampled individual (U)?*

Because the number of equations in the KFE model increases substantially for extra individuals or extra SNPs, we limited this calculation to a controlled system in which we explored all possible combinations of divergent SNPs ranging from 0 to 3. Moreover, we tested the effect of an increasing difference in time between the sampling of the sequences ($\Delta t = t^B - t^A$), from 0 to 10 years. The model (SEI, see main text Methods sections) and the parameters were chosen according to the pairs transmission calculation, and for the three parameters (infection rate, $\beta$, latency period, $1/\sigma$, and SNPs substitution rate, $\mu$) we used the median parameter obtained after the KFE calculation (respectively, 0.028 × contact × year, 1.29 years, and 0.16 base pair × genome × year).

As Figure S3.2 shows, in the case where two infected hosts *M. bovis* strain has no genetic difference (zero divergent SNPs in both), the probability of an intermediated infection with an unsampled individual was higher than for direct transmission when they are sampled more than four years apart. An increased number of divergent SNPs (both within the source and infected sequences) corresponded to a decreased temporal threshold. In fact, in the case where both A and B sequences included three divergent SNPs (total distance six), the three-individual transmission chain was more likely than the direct one for *Δt* of one year.

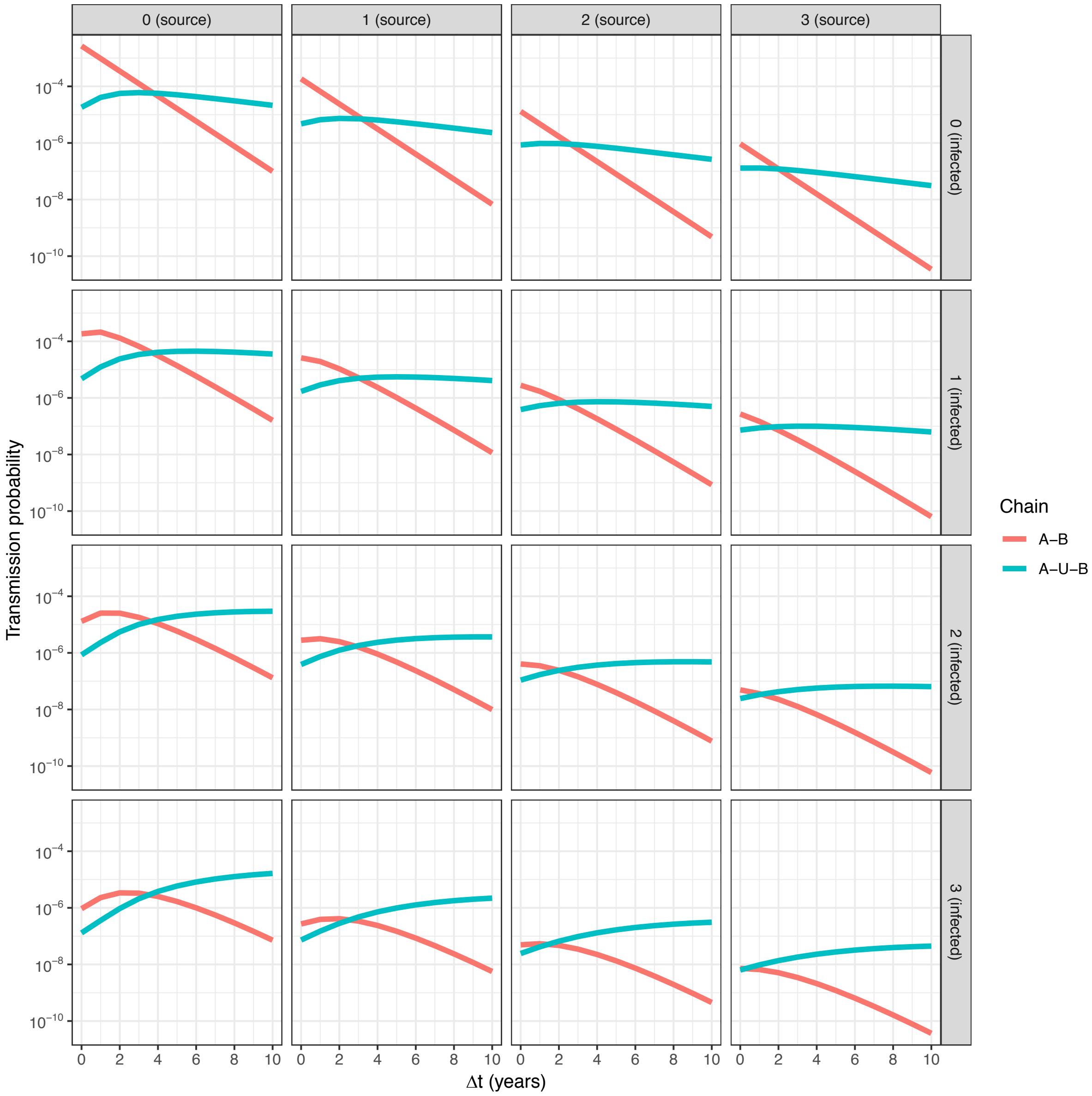


*Figure S3.2. Comparison of the transmission probability (y-axis) between two hosts with direct transmission (chain A-B, red line) and including an unsampled intermediary (chain A-U-B, blue line), for increasing time between the two hosts sampling (Δt). Each panel shows the result for a combination of source and infected sequence divergent SNPs, denoted in the labels.*
